# Tau Accumulation *via* Reduced BAG3-mediated Autophagy Is Required for GGGGCC Repeat Expansion-Induced Neurodegeneration

**DOI:** 10.1101/727008

**Authors:** Xue Wen, Ping An, Hexuan Li, Zijian Zhou, Yimin Sun, Jian Wang, Lixiang Ma, Boxun Lu

**Author notes:** These authors contributed equally to this work (Xue Wen, Ping An). Correspondence to: Boxun Lu, Lixiang Ma, Jian Wang.

## Abstract

Expansions of trinucleotide or hexanucleotide repeats lead to several neurodegenerative disorders including Huntington disease (HD, caused by the expanded CAG repeats (*CAGr*) in the *HTT* gene) and amyotrophic lateral sclerosis (ALS, could be caused by the expanded GGGGCC repeats (*G4C2r*) in the *C9ORF72* gene), of which the molecular mechanisms remain unclear. Here we demonstrate that loss of the *Drosophila* orthologue of tau protein (dtau) significantly rescued *in vivo* neurodegeneration, motor performance impairments, and shortened life-span in *Drosophila* models expressing mutant HTT protein with expanded *CAGr* or the expanded *G4C2r*. Importantly, expression of human tau (htau4R) restored the disease-relevant phenotypes that were mitigated by the loss of dtau, suggesting a conserved role of tau in neurodegeneration. We further discovered that *G4C2r* expression increased dtau accumulation, possibly due to reduced activity of BAG3-mediated autophagy. Our study reveals a conserved role of tau in *G4C2r*-induced neurotoxicity in *Drosophila* models, providing mechanistic insights and potential therapeutic targets.

## INTRODUCTION

An expansion mutation of the GGGGCC repeat (*G4C2*) within intron 1 of *C9orf72* is a common mutation associated with of sporadic amyotrophic lateral sclerosis (ALS) and frontotemporal dementia (FTD) (DeJesus-Hernandez et al., 2011; Renton et al., 2011). ALS is a fatal neurodegenerative disorder primarily affecting motor neurons, whereas FTD is causing neurodegeneration primarily in the frontal, insular, and anterior temporal cortex. The aggregation-prone dipeptides synthesized from the expanded *G4C2 via* repeat-associated non-ATG (RAN) translation and/or the *G4C2* RNA foci / membraneless granules originated *via* phase separation are likely the cause of neurodegeneration, but the downstream molecular mechanisms remain unclear. Abnormalities of the microtubule-binding protein tau play a central role in several neurodegenerative diseases termed tauopathies, including Alzheimer’s disease (AD), Progressive supranuclear palsy (PSP) and tau-positive FTD with parkinsonism (FTDP-17), etc.. Among them, FTDP-17 is a type of FTD caused by aberrant splicing of tau, and similar mechanisms mediate the pathology in Huntington disease (HD), another disease caused by nucleotide repeat expansion (the CAG repeat expansion in exon1 of the *HTT* gene) (Fernandez-Nogales et al., 2014). Thus, we investigated whether tau may mediate *G4C2*-induced neurotoxicity, which might share molecular commonalities in their pathological mechanisms with CAG repeat expansion diseases.

*Drosophila* models have been widely used to study genetic neurodegenerative disorders, especially CAG expansion and GGGGCC expansion diseases, probably because of their monogenetic nature. Many of the genetic or chemical modifiers identified in these models have been validated in patient cells and mammalian models. In this study, we first characterized the pathophysiological function of *Drosophila* orthologue of tau (dtau) in HD models. We confirmed a conserved role of dtau in HD neurotoxicity by showing that the loss of dtau significantly rescued phenotypes in HD *Drosophila* models expressing human mutant HTT protein (mHTT) exon1 fragment, consistent with the observation in an HD mouse model expressing the mHTT exon1 fragment (Fernandez-Nogales et al., 2014). We then investigated the potential role of dtau in *G4C2*-induced neurotoxicity in *Drosophila* models, and explored tau-relevant pathogenic mechanisms.

## RESULTS

### dtau knockout rescued HD-relevant phenotypes including *in vivo* neurodegeneration, behavioral deficits and shortened life-span in *Drosophila* models

In order to capture the neuronal morphology relatively clearly in the *Drosophila* brain *in vivo*, we utilized a simple *UAS-GAL4* system to express membrane GFP markers (mCD8GFP) in a very small group of neurons by *GMR61G12-GAL4*, so that their morphology could be clearly observed. We detected the *in vivo* HD-relevant neurodegeneration by observing the loss of major axon bundles in the GFP labeled by neurons induced by expression of human mHTT but not wtHTT exon1 fragment (*HTT.ex1.Q72* versus *HTT.ex1.Q25*, Fig. S1A). The neurodegeneration occurs in both dendrites and axons (Figure S2).

At the whole animal level, expressing mutant HTTexon1 (*HTT.ex1.Q72*) by the pan neural driver *elav-GAL4* caused motor performance impairments and shortened life-span compared to ones expressing the wild-type version (*HTT.ex1.Q25*, Figure S1B-C), recapitulating HD-relevant phenotypes in patients.

We then investigated the potential role of tau in HD by testing the knockout of the *Drosophila tau* gene to eliminate the expression of dtau. *Drosophila tau* is the homolog of human *MAPT*, the gene expressing the tau protein. The dtau knockout line we utilized was previously reported to present a specific deletion of the genetic region spanning from exon 2 to exon 6, which codes for the microtubule-binding region of dtau (Burnouf et al., 2016). Interestingly, loss of dtau significantly rescued HD-relevant *in vivo* neurodegeneration, behavioral deficits and shortened life-span phenotypes, suggesting that dtau mediates mHTT neurotoxicity, at least partially (Figure S1A-C). The observation was consistent with the study using the mouse R6/2 HD model, confirming a conserved role of dtau in neurodegeneration. Expressing human tau (htau4R) restored HD-relevant phenotypes in the dtau KO *Drosophila* (Figure S1C-D), further validating the conservation of dtau function in neurodegeneration.

### dtau knockout rescued expanded *G4C2*-induced ALS-relevant phenotypes including *in vivo* neurodegeneration, behavioral deficits and shortened life-span

The study using HD models demonstrated a conserved role of dtau in neurodegeneration, and thus we investigated potential functions of dtau in expanded *G4C2*-induced neurotoxicity in a well-characterized and widely used *Drosophila* model expressing 30 repeats of *G4C2* (*(G4C2)_30_*) (Xu et al., 2013). Similar as the HD *Drosophila* model, *UAS-mCD8GFP-GAL4* driven *(G4C2)_30_* expression led to *in vivo* neurodegeneration (Figure 1A). In addition, *(G4C2)_30_* expression in the motor neurons driven by *UAS-mCD8GFP-GAL4* led to significant reduction of neuro-muscular junctions (NMJs), which were detected by the NMJ marker Brp and axon marker HRP (Figure 1B). At the whole animal level, neuronal expression of *(G4C2)_30_* driven by *elav-GAL4* led to motor performance impairments and shortened life-span (Figure 1C-D). Loss of dtau significantly mitigated all these disease-relevant phenotypes (Figure 1A-D), suggesting that dtau may mediate neurotoxicity induced by expanded *G4C2* as well.

**Figure 1.**
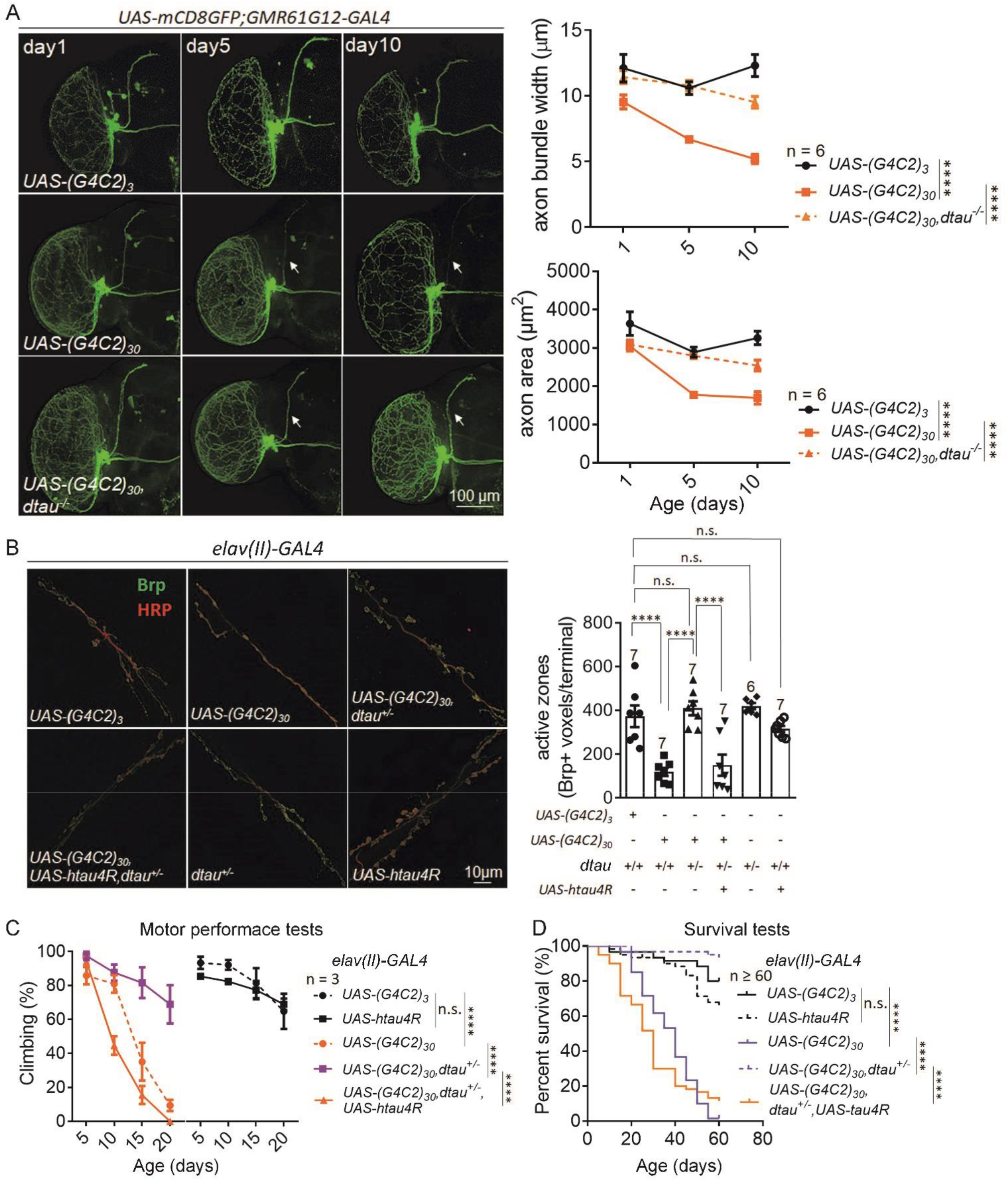
*dtau* knockout rescued *in vivo* neurodegeneration, motor function deficits, and shortened lifespan in *G4C2r* expressing flies. A) Similar as Figure 1A-B, but testing the expanded *G4C2r* induced toxicity. *dtau* KO significantly rescued *in vivo* neurodegeneration induced by *(G4C2)_30_* expression. B) Representative images and quantifications of neurodegeneration in the neuromuscular junctions (NMJs) in flies with the indicated genotypes. NMJs of muscle 6/7 in abdominal segment A2 and A3 of third instar larvae were stained with anti-HRP (neuronal axon marker, red channel) and anti-Bruchpilot (Brp) (active zone marker, green channel). The Brp-positive active zones were reduced by *(G4C2)_30_* expression, and rescued by *dtau* KO. The white arrows indicated the buttons of NMJs. *Scale* bars, 10 μm. C-D) Similar as Figure 1C-D, but for *G4C2* flies. *dtau* KO significantly rescued motor function deficits and shortened lifespan induced by *(G4C2)_30_* expression. For all plots in A-C, error bars indicate mean ± SEM. Asterisks denote statistically significant differences (*p≤0.05, **p≤0.01, ***p≤0.001, ****p≤0.0001).

We then examined whether dtau and human tau are conserved in mediating the expanded *G4C2*-induced toxicity by expressing htau4R in the neurons driven by *elav-GAL4*. htau4R expression restored *(G4C2)_30_* induced neurodegeneration, deficient motor performance and the shortened life-span phenotypes in the dtau KO *Drosophila* (Figure 1C-D), confirming that tau plays a conserved role in the expanded *G4C2*-induced neurodegeneration as well.

### The expanded *G4C2* increased tau protein levels in *Drosophila* and human cells

We then explored potential mechanisms regarding how dtau influenced the expanded *G4C2r*-induced toxicity. There are two major possibilities: 1. dtau is an upstream regulator of expanded *G4C2r* expression; 2. dtau is a downstream factor that is modulated by expanded *G4C2r* and mediates its neurotoxicity. Thus, we first examined whether dtau knockout lowered the expanded *G4C2r* RNA, which is likely the fundamental source of neurotoxicity in this model (Xu et al., 2013). Since it is extremely difficult to amplify the *G4C2* repeat region *per se* by quantitative PCR, we chose the upstream and downstream sequences for qPCR measurements (Figure S3A). We observed no significant change in expanded *G4C2r* mRNA levels in the dtau heterozygous knockout flies, suggesting that dtau is not an upstream regulator of expanded *G4C2r* expression (Figure S3B).

The evidence above suggested that dtau is more likely a downstream mediator of the expanded *G4C2r*-induced toxicity. We first examined potential missplicing of tau transcripts. Alternative splicing of exon 10 of *MAPT* gene results in tau isoforms containing either three or four microtubule-binding repeats (3R and 4R, respectively) (Liu and Gong, 2008). Similar as the human tau, dtau proteins expressed from different *tau* transcripts (Figure S3C) contain either three or four putative tubulin-binding repeats (MTBDs) that are homologous with mammalian MTBDs (Heidary and Fortini, 2001). In addition, all dtau proteins have an extra MTBD that is homologous with *Caenorhabditis elegans* but not mammalian ones. Thus, dtau have the 4R versus 3R isoforms due to alternative splicing (or 5R versus 4R if we consider the extra MTBD), corresponding to 4R versus 3R isoforms in human tau. We thus measured different *dtau* transcript levels and the 3R/4R ratio in fly brains with *(G4C2)_30_* expression versus the control (no repeat or *(G4C2)_3_*), and observed no significant changes in either the transcript levels or the 4R/3R ratio (Figure S3D), suggesting that the expanded *G4C2r* may not induce tau missplicing.

Increased level of tau protein is another possible driver of neurodegeneration. We thus investigated whether tau protein level is increased in the expanded *G4C2r* expressing *Drosophila* model.

Expression of expanded *G4C2r* (*(G4C2)_30_*) led to increased tau protein levels in tissue samples from 3^rd^ stage larvae, pupae and adult *Drosophila* (Figure 2A-C), suggesting that the expanded *G4C2r* expression increases tau accumulation. Consistent with the observation in *Drosophila*, exogenous expression of the expanded *G4C2r* (*(G4C2)_25_*) in HEK293T cells also resulted in tau protein accumulation (Figure 2D). We further investigated levels of tau aggregates, which is likely a toxic biomarker associated with tau accumulation. Consistently, HeLa cells with expanded *G4C2r* expression showed drastic increase of tau aggregates (Figure 2E), suggesting that expanded *G4C2r* is capable of enhancing aggregated form of tau as well.

**Figure 2.**
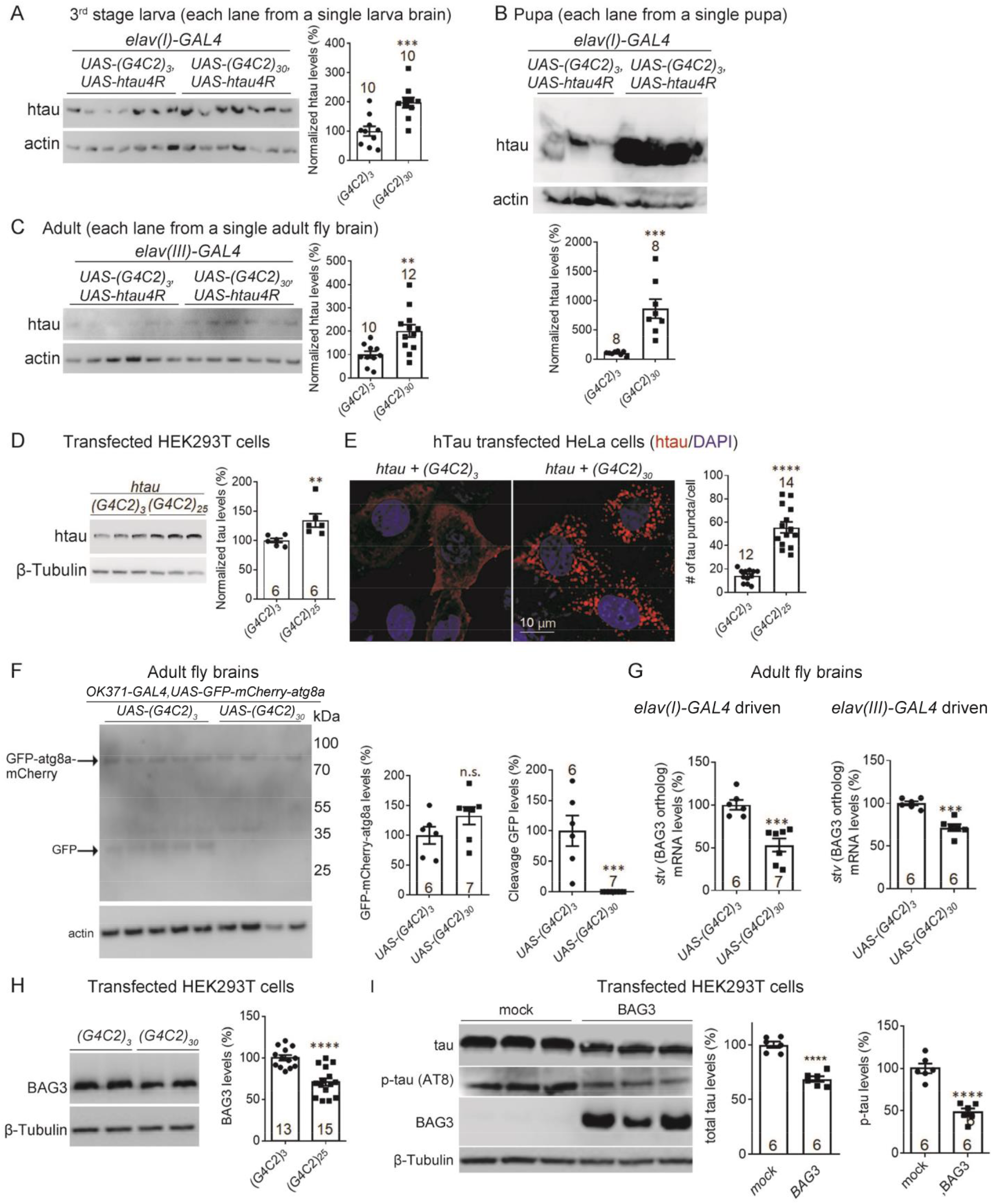
*G4C2r* expression increased tau protein levels *in vivo* and in cells *via* reducing BAG3-mediated autophagy. A-C) Representative Western-blots and quantifications showing that the total tau levels were increased by *(G4C2)_30_* expression in the 3rd instar larva, pupa and adult stage flies, compared to the *(G4C2)_3_* expression controls. n indicates the number of individual flies. The statistical analysis was performed by two-tailed unpaired t tests. D) Representative immunofluorescence images of whole-mount fly brains showing increased transgenic htau levels in *(G4C2)_30_* flies. All the flies was dissected and stained at day 16. E) Representative Western-blots and quantifications showing that total tau levels were increased by *(G4C2)_25_* compared to *(G4C2)_3_* controls in transfected HEK293T cells. The statistical analysis was performed by two-tailed unpaired t tests. F) Representative immunofluorescence images and quantifications (measured by the number of tau puncta *per* cell) showing that *(G4C2)_25_* expression increased tau aggregates in HeLa cells, compared to the *(G4C2)_3_* expressing controls. The statistical analysis was performed by two-tailed unpaired t tests. The aggregates were tested in HeLa cells, because tau aggregates were hardly visible in HEK293T cells. G) Representative Western-blots and quantifications showing that the cleaved product (free GFP cleaved from GFP-mCherry-atg8) band is largely missing in motor neurons expressing *(G4C2)_30_*, suggesting an inhibition of autophagy activity in these neurons. n indicates the number of individual flies. The statistical analysis was performed by two-tailed unpaired t tests. H) qPCR showing a reduction of *starvin* (*stv*) mRNA levels by *(G4C2)_30_* expression compared to the *(G4C2)_3_* expressing controls. The results were consistent in both *elav(I)-GAL4* or *elav(III)-GAL4* driven flies. I) Representative Western-blots and quantifications of lysates from transfected HEK293T cells showing that BAG3 protein levels were reduced by *(G4C2)_25_* expression compared to the *(G4C2)_3_* expressing controls. For all plots, bars indicate mean ± SEM. Asterisks denote statistically significant differences (*p≤0.05, **p≤0.01, ***p≤0.001, ****p≤0.0001).

We further investigated the levels of phosphorylated tau (p-tau), which is considered as a main toxic species that causes neurodegenerations. Western-blots of brains of adult *Drosophila* expressing expanded *G4C2r* showed an increase of certain forms of phosphorylated tau (p-tau) as detected by the p-tau antibody ps262 or AT8 (Figure S4A). Consistently, the p-tau increase by expanded *G4C2r* expression was observed in HEK293T cells as well (Figure S4B). The p-tau increase is likely a subsequence of increase tau accumulation (Figure 2A-E), and may also contribute to the neurotoxicity induced by the expanded *G4C2r* expression.

### Expanded *G4C2r* expression decreased *BAG3* (*starvin*) mRNA levels and impaired the autophagy flux

The tau protein is known to be degraded *via* autophagy in the cells (Zhu et al., 2017), and thus we hypothesized that expanded *G4C2r* expression impaired autophagy function, resulting in tau accumulation as a consequence. We assayed autophagy flux *in vivo* by testing the cleavage of GFP-mCherry-atg8a. atg8a (autophagy-related protein 8a) in *Drosophila* is the homolog of the mammalian protein LC3, which is the key autophagosome protein that gets cleaved in lysosomes. Thus, GFP-atg8a or GFP-LC3 cleavage is a widely used assay for measurements of the autophagy flux (Kimura et al., 2007). By expressing *GFP-mCherry-atg8a* driven by *OK371-GAL4* in *Drosophila*, the autophagy flux in the motor neurons (expressing *OK371*) could be measured by the levels of free GFP, the cleaved product of the GFP-mCherry-atg8a protein. The free GFP band essentially disappeared in all the *(G4C2)_30_* expressing *Drosophila* brains tested compared to the ones from *(G4C2)_3_* expressing brains (Figure 2F), indicating a drastic impairment of autophagy flux. Consistently, the two other functional readout for autophagy flux including the levels of ubiquitinated proteins in the lysate precipitations and the ref(2)P protein (the *Drosophila* orthologue of *SQSTM1/p62* in mammals (DeVorkin and Gorski, 2014)) were significantly increased by *(G4C2)_30_* expression compared to the *(G4C2)_3_* expressing or the wild-type (*w1118*) controls (Figure S5), further demonstrating that the autophagy flux was impaired by the expanded *G4C2r* expression.

We then explored the potential molecular mechanisms of autophagy impairment. Studies have found that BAG3 (BAG family molecular chaperone regulator 3), a key autophagy modulator (Kathage et al., 2017), is negatively associated with tau protein levels (Ji et al., 2019; Lei et al., 2015). We thus hypothesized that the expanded *G4C2r* expression may reduce BAG3 levels, leading to impaired autophagy function and tau accumulation. Consistent with this hypothesis, *(G4C2)_30_* expression in *Drosophila* significantly reduced the mRNA levels of *starvin* (*stv*), the *Drosophila* orthologue of BAG3 (Arndt et al., 2010), compared to the *(G4C2)_3_*-expressing controls (Figure 2G). Due to a lack of anti-starvin antibody, we examined potential BAG3 protein level changes in HEK293T cells, and observed consistent BAG3 lowering by the expanded *G4C2r* expression (Figure 2H). To further confirm that BAG3 may functionally modulate tau protein levels, we knocked-down BAG3 in HEK293T cells co-expressing *(G4C2)_25_* and tau, and observed increased the tau level by lowering BAG3 expression (Figure S6A). Consistently, overexpressing BAG3 reduced tau levels (Figure 2I). Taken together, the expanded *G4C2r* expression decreased BAG3 levels, leading to impaired autophagy function and increased tau levels, or at least making a partial contribution of tau increase.

## DISCUSSION

Taken together, our study revealed a functional role of tau in *G4C2* mediated neurotoxicity. While some of the previous studies suggest a potential link between tau and *G4C2* expansion, there has been no study demonstrating tau as a potential mediator of neurotoxicity induced by *G4C2* expansion. Thus, we have made a novel discovery that tau may mediated the expanded *G4C2r*-induced neurotoxicity by showing that knockout of dtau mitigated *in vivo* neurodegeneration, motor performance deficits and shortened lifespan in the *Drosophila G4C2* model (Figure 2).

While our study was mainly performed in the *Drosophila* model, tau’s function in *G4C2*-induced neurotoxicity is likely conserved between human and flies. dtau has previously been cloned and demonstrated to displays microtubule-binding properties (Heidary and Fortini, 2001), but its potential role in neurodegeneration has been controversial. Studies using either *Drosophila* RNAi lines or tau hypomorphic/deficiency lines led to contradictory results regarding the potential detrimental or beneficial effects of dtau removal, possibly because of off-target effects or incomplete removal of dtau, and there was no disease phenotype “restoration” experiments expressing human tau (htau) to test the functional similarity between dtau and htau. Meanwhile, dtau knockout did not trigger major effects in terms of fly survival and fly climbing ability (Figure S1 & Figure 1), and it did not influence the toxic effects on fly survival related to the expression of human Aβ, suggesting that dtau may mediate neurodegeneration in certain context, and not in a nonspecific manner. On the other hand, our study demonstrated that expression of htau restored most phenotypes that were rescued by the dtau knockout (Figure S1C-D & 1B-D). In addition, the expanded *G4C2r* expression increased tau levels in both *Drosophila* and mammalian cells (Figure 2A-E). Finally, dtau knockout rescued HD-relevant phenotypes in *Drosophila* expressing mHTT proteins (Figure S1), consistent with the observation in a mouse model (Fernandez-Nogales et al., 2014). These pieces of cross-species evidence suggest a conserved role of dtau in mediating neurodegeneration, and justify using dtau for mechanistic studies of tauopathies, providing a much faster and more convenient model for research in this field compared to mouse models.

BAG3-mediated autophagy may play a key role in maintaining cellular homeostasis under stress as well as during aging (reviewed by (Behl, 2016)), when the increased demand for protein degradation enhances BAG3 expression, followed by a functional switch from BAG1-mediated proteasomal to BAG3-mediated autophagic pathway. Meanwhile, potential changes or functional roles of BAG3 in *G4C2* models was previously unknown, and our study revealed a potential role of the conserved BAG3-autophagy pathway in the expanded *G4C2r*-induced neurotoxicity.

In summary, our study revealed an evolutionarily conserved BAG3-mediated autophagy-dependent mechanism between the expanded *G4C2r*-induced ALS and tauopathies, providing entry points for targeting these diseases.

## ACKNOWLEDGMENTS

We’d like to thank Prof. Peng Jin and Ranhui Duan for providing the *G4C2 Drosophila* models. Prof. Partridge L, Haihuai He and Peng Lei for providing *dtau* knockout *Drosophila* models. Prof. Yongqin Zhang for providing *UAS-htau4R, elav(I)-GAL4*, and related tool *Drosophila* stocks. We’d also like to thank National Natural Science Foundation of China (81870990, 91649105), National Key Research and Development of Program of China (2016YFC0905101), and Hereditary Disease Foundation for funding supports.

## SUPPLEMENTARY FIGURES

**Figure S1.**
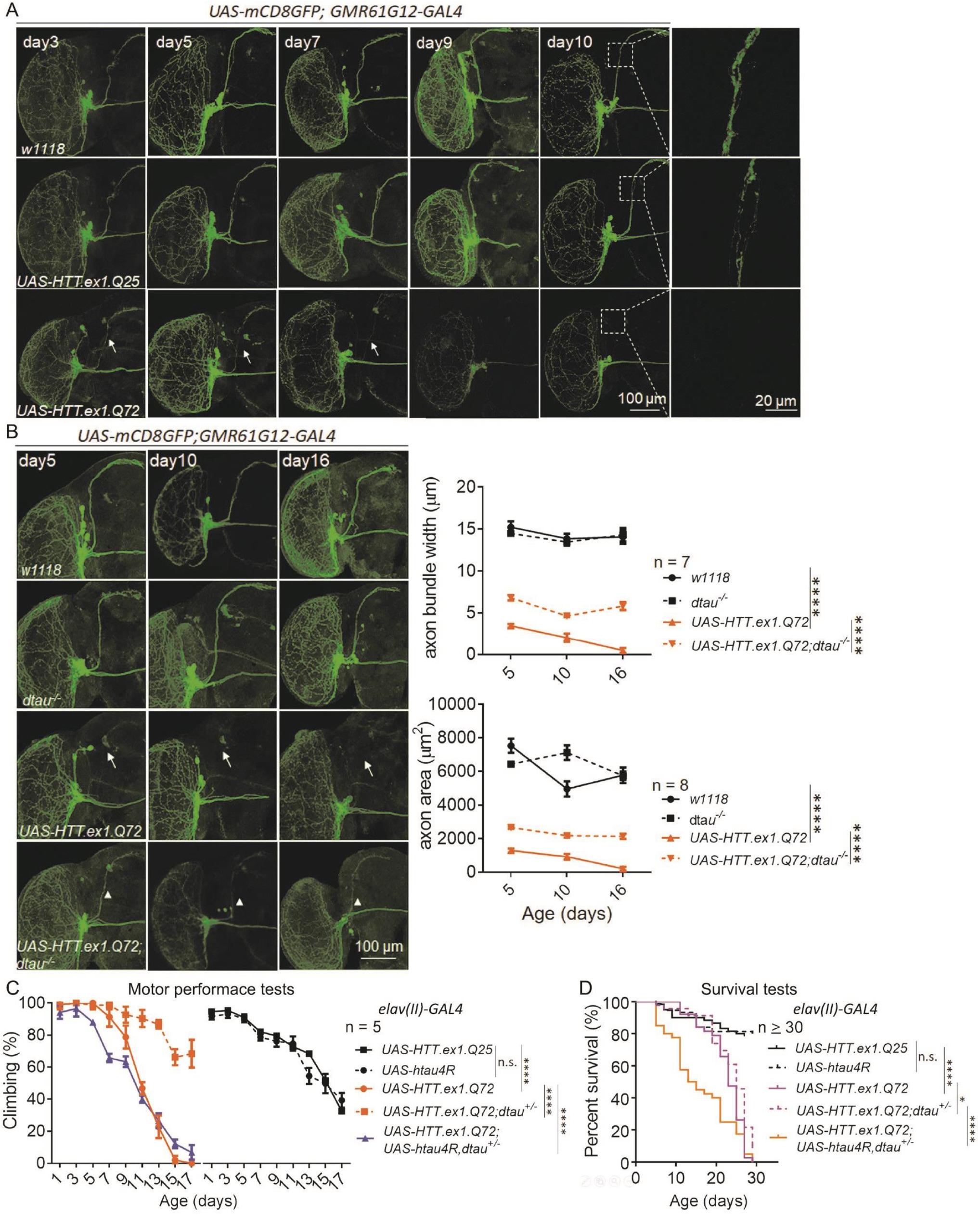
*dtau* knockout rescued *in vivo* neurodegeneration, motor function deficits, and shortened lifespan in HD flies. A) Representative immunofluorescence images of whole-mount brains of flies of indicated genotypes at indicated ages showing *in vivo* neurodegeneration. The morphology of small ventral lateral (sLNv) clock neurons was labeled by mCD8GFP protein, which was driven by *GMR61G12-GAL4*. The HTT.ex1.Q72 or HTT.ex1.Q25 expression was also driven by the same *GAL4*. The white arrows indicated a major axon bundle that were degenerated in the HTT.ex1.Q72 expressing flies, compared to the HTT.ex1.Q25 expressing or the w1118 control flies. *Scale* bars, 100 μm and 20 μm. B) Similar as A, but with or without *dtau* knockout (KO) as indicated. *dtau* Ko significantly rescued the *in vivo* neurodegeneration phenotype in the sLNv neurons. The plots (mean and SEM) on the right exhibit the quantification of axon bundle width and total axon area based on the GFP images. n indicates the number of individual flies. The statistical analysis was performed by two-way ANOVA and Dunnett’s post-hoc test. *Scale* bars, 100 μm. *dtau* KO significantly rescued *in vivo* neurodegeneration induced by mHTT. C) Motor performance tests measuring climbing ability of flies with indicated genotypes and ages. *dtau* KO significantly rescued motor function deficits induced by mHTT. n indicates the number of independently tested vials, which contained 15 virgin female flies in each one. The statistical analysis was performed by two-way ANOVA and Dunnett’s post-hoc test. D) Lifespan measurements of flies with indicated genotypes. *dtau* KO significantly rescued the shortened life-span induced by mHTT. n indicates the number of individual flies. The statistical analysis was performed by log-rank tests. For all plots in A-C, error bars indicate mean ± SEM. Asterisks denote statistically significant differences (*p≤0.05, **p≤0.01, ***p≤0.001, ****p≤0.0001).

**Figure S2.**
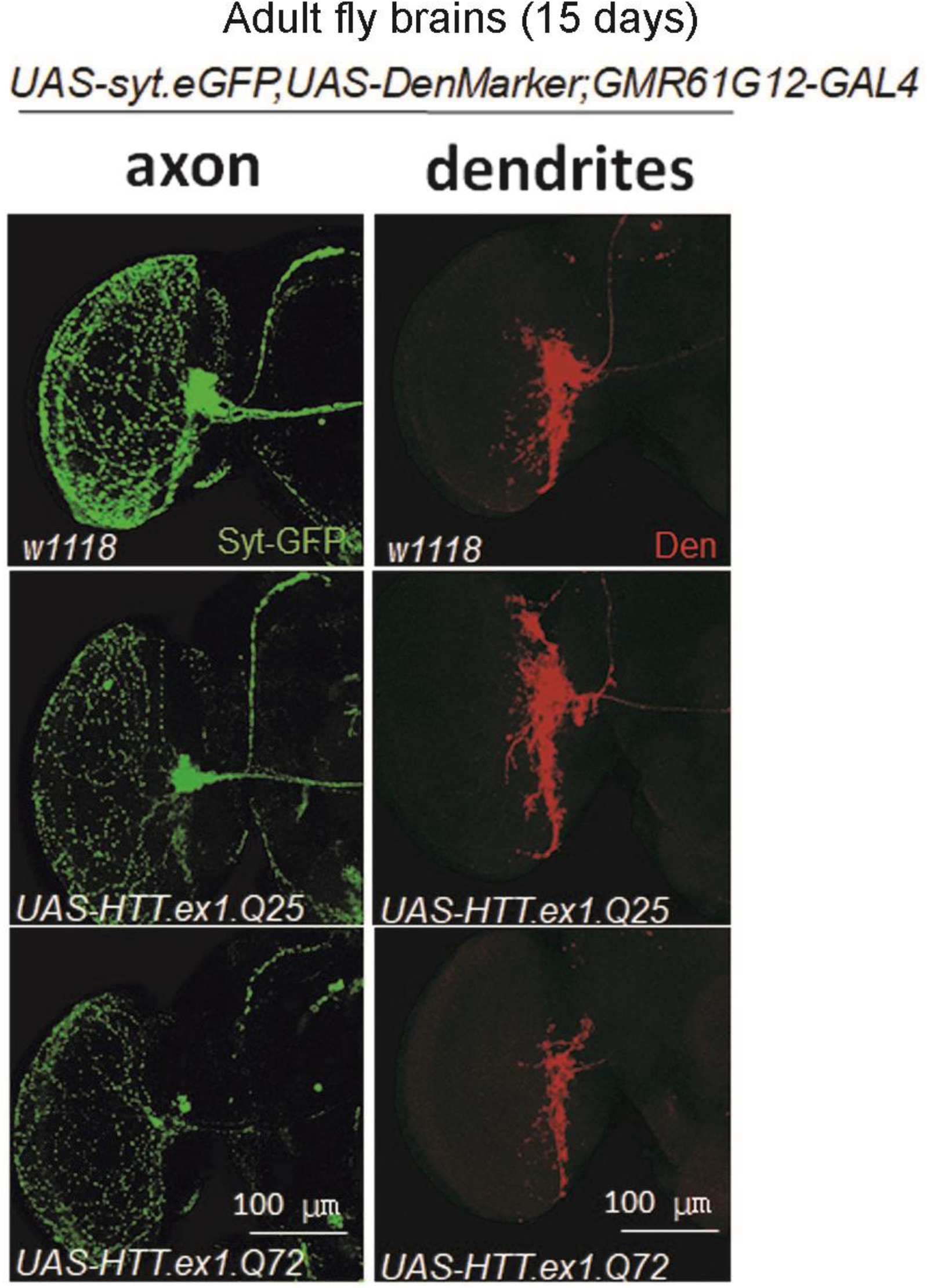
Validation of neurodegeneration detection by axon and dendrite markers. Representative immunofluorescence images of whole-mount brains from *Drosophila* of indicated genotypes. The expression of axon marker (syt.eGFP) or the dendrite marker (DenMarker) were driven by *GMR61G12-GAL4* in the sLNv clock neurons with HTT.ex1.Q25 or HTT.ex1.Q72 expression, or in the wild-type background (w1118). Obvious neurodegeneration could be detected in both axons and dendrites by HTT.ex1.Q72 expression.

**Figure S3.**
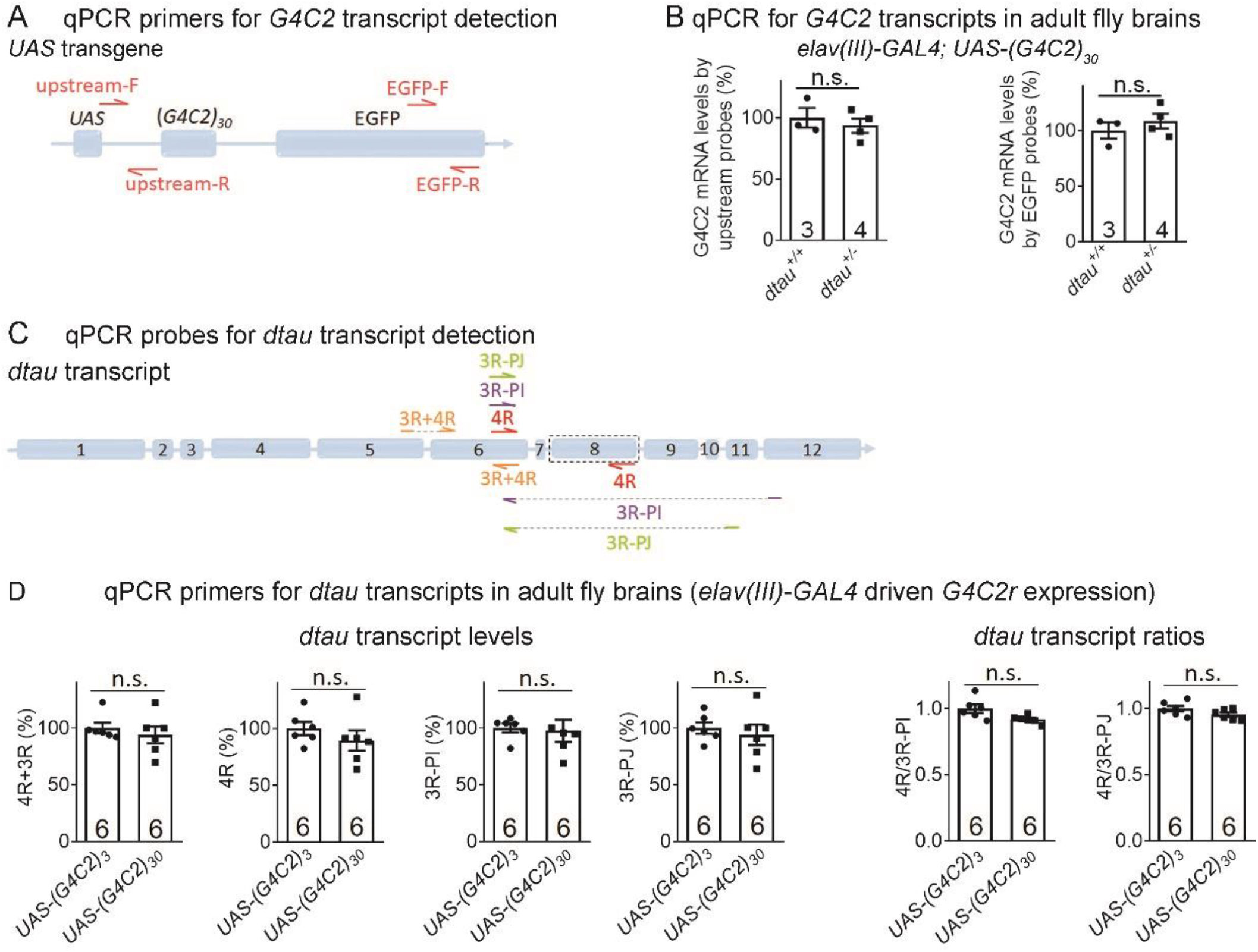
dtau and the expanded *G4C2r* did not influence each other’s mRNA levels. A) The schematic picture of the qPCR primers for measurements of the *(G4C2)_30_* transcript level. B) qPCR measurements of *(G4C2)_30_* transcripts in brain tissues from flies of indicated genotypes using indicated qPCR primers. n indicates the number of individual flies. The statistical analysis was performed by two-tailed unpaired t tests. C) The schematic picture of the qPCR primers for measurements of the endogenous *dtau* transcript level. D) qPCR measurements of the indicated *dtau* transcripts in brain tissues from flies of indicated genotypes. The statistical analysis was performed by two-tailed unpaired t tests. Neither the dtau transcript levels nor their splicing were influenced by *(G4C2)_30_* compared to *(G4C2)_3_*. For all plots, bars indicate mean ± SEM. Asterisks denote statistically significant differences (*p≤0.05, **p≤0.01, ***p≤0.001, ****p≤0.0001). For all the qPCR primers tested, the no reverse transcriptase control showed no amplification, validating that the signals were not contaminated by genomic DNA.

**Figure S4.**
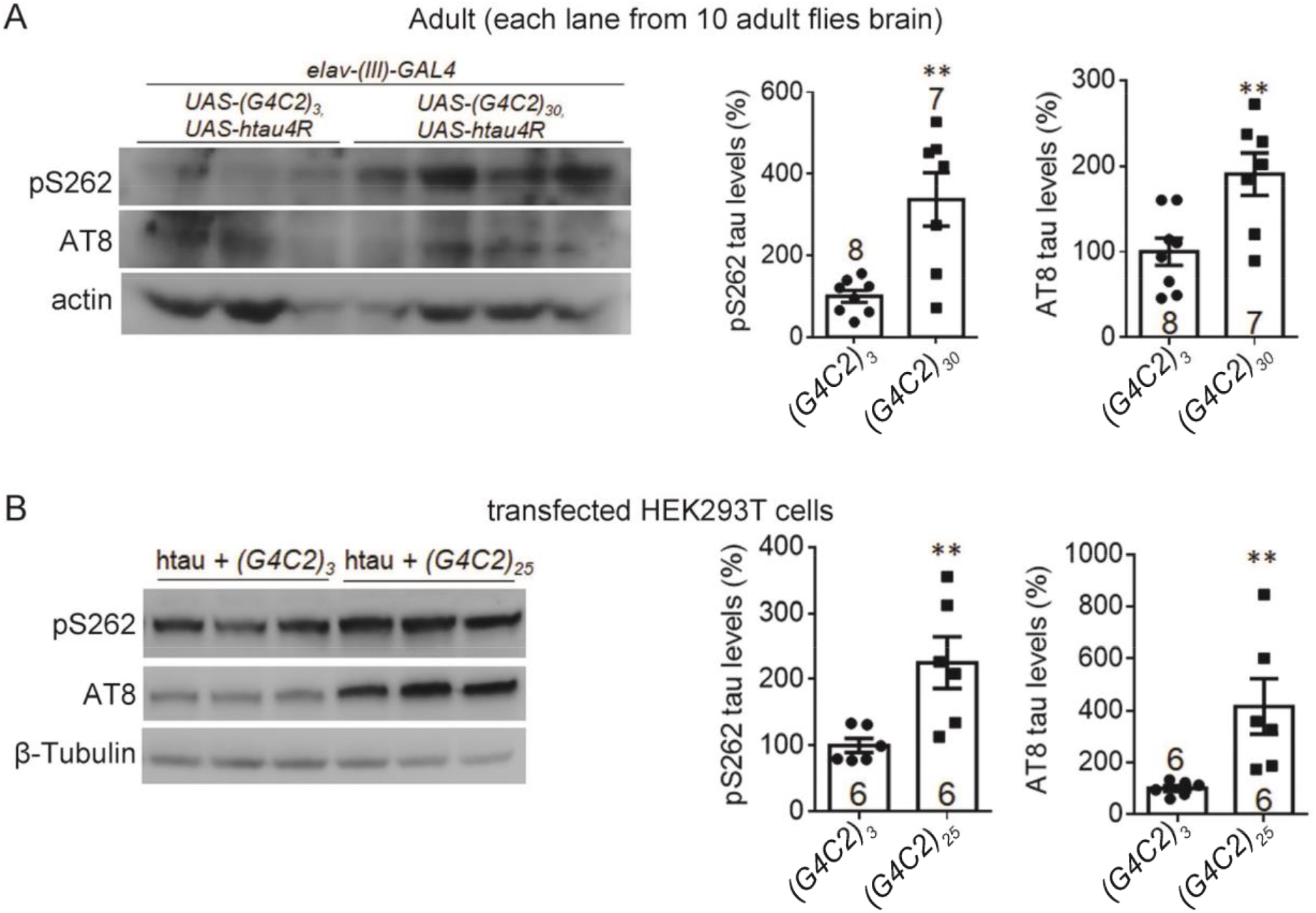
*G4C2r* expression increased certain forms of phosphorylated tau protein levels *in vivo* and in cells. A) Representative Western-blots and quantifications showing that the phosphorylated tau levels detected by the antibody pS262 or AT8 were increased by *(G4C2)_30_* expression in the adult fly heads, compared to the *(G4C2)_3_* expressing controls. n indicates the number of biological replicates. The statistical analysis was performed by two-tailed unpaired t tests. B) Similar as A, but in transfected HEK293T cells.

**Figure S5.**
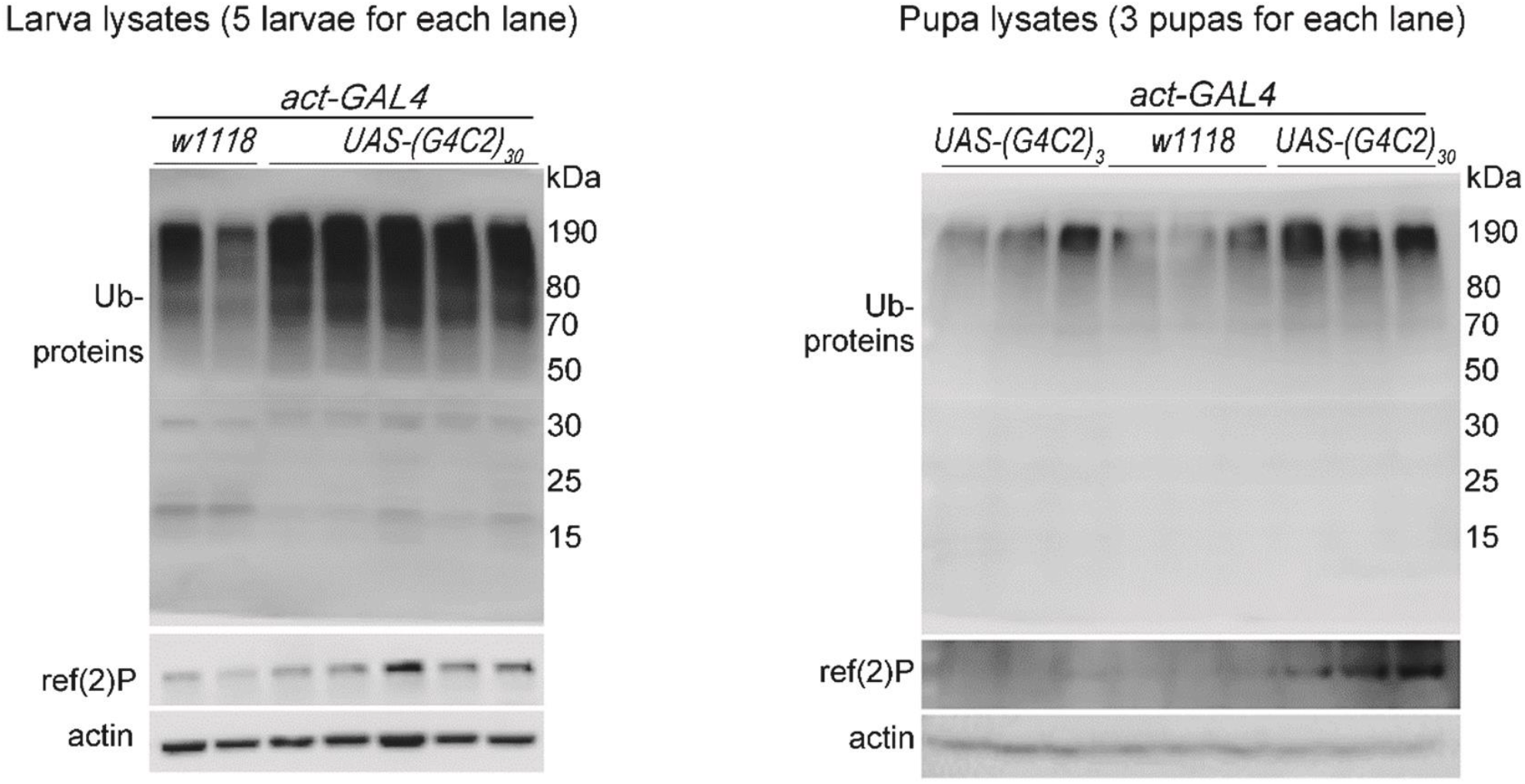
*G4C2r* expression reduced autophagy flux. Representative Western-blots of ubiquitinated proteins showing an increase of these proteins in *(G4C2)_30_* expressing larva or pupae.

**Figure S6.**
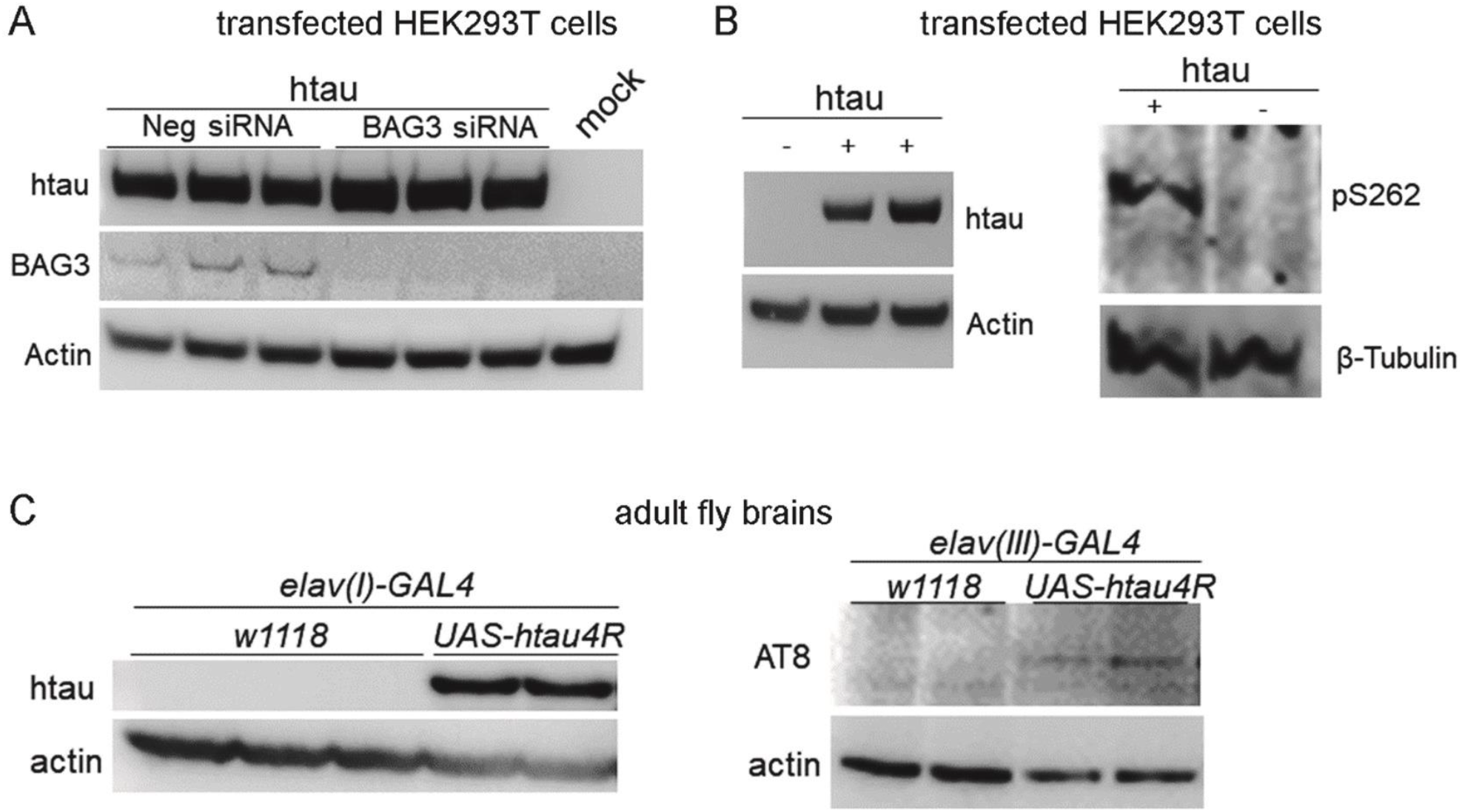
Validation of specificity of major antibodies. A-C) Representative Western-blots of HEK293T and fly samples showing the specificity of anti-total tau and phos-tau, including htau, pS262 and AT8 antibodies. The representative Western-blots in A) also showed that BAG3 knock-down increased tau (detected by the antibody 5A6) levels.

## MATERIAL AND METHODS

### Fly stocks and genetics

Fly culture and crosses were performed on standard food according to standard procedures and raised at 25 °C. The *elav(c115)-GAL4* (*elav(I)-GAL4*) (no.458), *elav(II)-GAL4* (no.8765), *elav(III)-GAL4* (no.8760), *OK371-GAL4* (no.26160) and *UAS-GFP-mCherry-atg8a* (no.37749) were obtained from the Bloomington Drosophila Stock Center at Indiana University (Bloomington, IN, USA). The *dtau^−/−^* stock was described previously (Burnouf et al., 2016).*GMR61G12-GAL4* was obtained from the FlyLight *GAL4* collection organization (http://flweb.janelia.org/cgi-bin/flew.cgi). *UAS-(G4C2)_30_-EGFP* and *UAS-(G4C2)_3_-EGFP* lines used in this study were described previously (Xu et al., 2013). The transgenic *Drosophila* lines expressing human wild-type was described previously (Wittmann et al., 2001). The expression of *(G4C2)_30_* in all cells of the peripheral and central nervous system using *elav(I)-GAL4* caused lethality in early development as described previously (Xu et al., 2013). As the pupal lethality of *(G4C2)_30_* driven by *elav(I)-GAL4* precluded studies in mature neurons of the adult brain, and thus we used *elav(II)-GAL4* or *elav(III)-GAL4* for substituting it and overcome the shortage. *UAS-HTT.ex1.Q25* and *UAS-HTT.ex1.Q72* were generated by injecting *pUAST-HTT.ex1.Q25* or *pUAST-HTT.ex1.Q72* plasmid with helper-plasmids named *Δ2-3* into *w1118*.

### Behavioral and lifespan experiments

For behavioral experiments (climbing assay), we placed 15 age-matched virgin female flies in an empty vial and tapped them down. The percentage of flies that climbed past a 9-cm-high line after 15 s was recorded. The mean of five observations is plotted for each vial on each day, and data from multiple vials containing different batches of flies were plotted and analyzed by two-way ANOVA tests. The flies were randomly placed into each tube. For lifespan measurements, we placed ⩾60 age-matched virgin female flies in an empty plastic vial and recorded the survival situation for each vial on each day. For both behavioral and lifespan measurement experiments, the person who performed the experiments were blind to the genotypes until data analysis. The survival distribution of the two genotypic groups were compared using the log-rank (Mantel-Cox) test.

### Plasmids used for fly strain generation and mammalian cell transfection

The HTT.ex1.Q25 and HTT.ex1.Q72 cDNA were achieved by PCR amplification from pcDNA-HTT.ex1.Q25 and pcDNA-HTT.ex1.Q72 plasmid, and they were cloned into pUAST vector. The tau-snap plasmid were generated by inserting the cDNA expressing one transcript of human tau (0N4R, cloned by PCR amplification from the UAS-htau4R Drosophila strain) into the snap-tag vector (NEB, cat. no. P9312S). (G4C2)_3_ and (G4C2)_25_ plasmids were generated by inserting (g4c2)_3_ or (g4c2)_25_ repeats into pTT-sfGFP vector. The (g4c2)_3_ repeats was achieved by primer annealing process and (g4c2)_25_ repeats was achieved with using the enzyme to cut the plasmid named pHR-Tre3G-29xGGGGCC-12xMS2 (addgene, cat.no. #99149). The BAG3 vector was constructed by cloning human BAG3 cDNA sequence from PUC57_BAG3wt-eGFP (addgene,cat.no. #98182) into pTT-sfGFP vector. As the end of PCR product BAG3 gene was with termination codon, GFP protein in the vector was not expressed with BAG3 protein.

### Immunostaining

For immunofluorescence of cultured HeLa cells (Figure 2E, cells were fixed in 4% paraformaldehyde (PFA) at room temperature for 10 min after washing with 1 × PBS for three times (20 minutes each), and then washing and permeabilized in 0.5% (vol/vol) TritonX-100 for 10 min. The cells were then blocked in blocking buffer (4% BSA + 0.1% (vol/vol) Triton X-100 in 1 × PBS) for 30 min and incubated overnight at 4 °C with the primary antibody (anti-tau antibody, DSHB, 5A6), and then washed three times with blocking buffer and incubated with secondary antibody at room temperature for 1 h. Coverslips were then washed three times, stained with 0.5 mg/ml DAPI for 5 minutes at room temperature, and then mounted in vectashield mounting medium (Vector, cat.no. H-1002). Images were taken by Zeiss Axio Vert A1 confocal microscopes and analyzed blindly by ImageJ for htau puncta numbers in each cell.

For imaging of in vivo neurodegeneration in the fly brains (Figure S1A-B, Figure 1A & Figure S2), the whole brains of adult flies at indicated ages were dissected on ice and then fixed by 4% PFA on ice for 20 min. The immunostaining was then performed in the same way as in cultured HeLa cells. The primary antibodies used were anti-GFP antibody (ProteinTech, cat. no. 50430-2-AP). The red fluorescence signals of DenMarker (Figure S2) were imaged directly without antibody staining. For NMJ immunostaining, the 3^rd^ larval stage flies were used for dissection to isolate the muscle tissues. The dissected samples were then fixed in 4% PFA at room temperature for 30 min, and then wash them by 0.5% PBST 3 times, each time about 20 min. The immunostaining was then performed similarly as in cells. The primary anti-Brp antibody nc82 (DSHB, cat. no. AB 2314866) were then added to the samples at 1:20 for incubation at 4 °C overnight. The samples were then washed for 3 times (20 minutes each), then incubated with the fluorophore-labeled primary anti-HRP antibody Cy3-HRP (Jackson, cat. no. 123-165-021; diluted at 1:200) and the secondary antibody (goat anti-mouse, 633nm) for 1 hour at room temperature. The samples were then washed three times and mounted in vectashield mounting medium (Vector, cat.no. H-1002). Images were taken by Zeiss Axio Vert A1 confocal microscopes and analyzed blindly by ImageJ. The NMJ locating at muscle 6/7 of segment A2 or A3 were chosen for analysis.

### Western-blot experiments and antibodies

For protein extraction from the fly tissues, the samples (brains or whole bodies) were dissected on ice and homogenized with a tissue grinder for 5 min at 60 Hz and lysed on ice for 60 min in the lysis (1× RIPA buffer (Beyotime, cat. no. #P0013B) + 1 × Complete protease inhibitor (Roche, cat. no. 4693159001)). The samples were then sonicated for 10 cycles, 15 s on and 20 s off, and then collected. The whole lysates were then loaded onto the 4-12% bis-tris gradient gel for Western-blots.

For the Western-blots of ubiquitinated proteins and ref(2)P (Figure S5), the lysates were centrifuged at > 20,000g at 4°C for 30 min. The precipitates were then collected and loaded for Western-blots. For protein extraction from cells, the cell pellets were collected and lysed on ice for 30 minutes in 2% SDS (in 1× PBS + 1× Complete protease inhibitor (Roche, cat. no. 4693159001)) and sonicated for 10 cycles, 15 s on and 20 s off. The whole lysates were then collected and loaded for Western-blots.

### Antibodies used for Western-blots and immunostaining

anti-Brp (DSHB, cat. no. AB 2314866,); anti-htau (clone 5A6, detecting total human tau, DSHB, cat. no. AB 528487); anti-Actin (Millipore, cat. no. 92590); anti-β-Tubulin (Abcam, cat.no. #ab6046); anti-ref(2)P (Abcam, cat. no. #ab178440); anti-GFP (ProteinTech, cat. no. 50430-2-AP); anti-BAG3 (ProteinTech, cat. no. 10599-1-AP,); anti-ubiquitin (Dako, cat. no. Z0458); AT8 (detecting phosphorylated-tau, ThermoFisher, cat.no. MN1020); pS262 (detecting Ser262 phosphorylated-tau, ThermoFisher, cat. no. OPA1-03142); pS199 (detect Ser199 phosphorylated-tau, ThermoFisher, #701054). All the antibodies’ specificity has been validated in this study (Figure S6) or in previous literature (such as cited or indexed in Antibodypedia).

### RNA extraction and RT-qPCR

RNA from fly tissues or cells was extracted using RNAprep Kit (Tiangen, #DP419) followed by purification using the RNA-clean kit (Tiangen, #DP412) to remove proteins and RNase-free DNase I (Tiangen, #RT411) treatment to break down the genomic DNA. cDNA was obtained by reverse transcription using the FastQuant RT Kit with the oligo (dT) primer (Takara, #RR047A). qPCR was then performed using SYBR Green Realtime PCR Master Mix (Toyobo, #QPK-201). All the primers were published and validated previously, and further tested for standard curve and melting curve. Amplification efficiency was between 95%-105% and the R^2^ for linear relationship is > 0.999 for all primers. No reverse-transcriptase controls were used to ensure the specificity of the signals. qPCR primer sequences were as follows:

3R+4R-F 5’ggcggcgagaagaagata3’

3R+4R-R 5’gcgaaccgattttggactt3’

4R-F 5’tgggctcgacggccaatgtgaaaca3’

4R-R 5’ccgccaccgggcttgtgctttaca3’

3R-PI-F 5’aaggacaaggccaagccgaaggtg3’

3R-PI-R 5’ggtgccttccaatacttgatgtctccgc3’

3R-PJ-F 5’aaggacaaggccaagccgaaggtg3’

3R-PJ-R 5’tggactcttgatgtctccgccaccc3’

EGFP-F 5’tatatcatggccgacaagca3’

EGFP-R 5’gttgtggcggatcttgaagt3’

Upstream-F 5’tcaattaaaagtaaccagcaacca3’

Upstream-R 5’tccctattcagagttctcttcttgta3’

### Statistical analysis

Statistical comparisons between two groups were conducted by the unpaired two-tailed t tests. Statistical comparisons among multiple groups were conducted by one-way ANOVA tests and post-hoc tests for the indicated comparisons (Dunnett’s tests for comparison with a single control, and Bonferroni’s tests for comparison among different groups). Statistical comparisons for serials of data collected at different time points were conducted by two-way ANOVA tests. The similarity of variances between groups to be compared was tested when performing statistics in GraphPad Prism 7 and Microsoft Excel 2016. Normality of data sets was assumed for ANOVA and t tests, and was tested by Shapiro-Wilk tests. When the data were significantly different from normal distribution, nonparametric tests were used for statistical analysis. All statistical tests were unpaired and two-tailed.

